# Bidirectional locomotion induces unilateral limb adaptations

**DOI:** 10.1101/2024.08.22.609228

**Authors:** Russell L. Hardesty, Helia Motjabavi, Darren E. Gemoets, Jonathan R. Wolpaw

## Abstract

Humans can acquire and maintain motor skills throughout their lives through motor learning. Motor learning and skill acquisition are essential for rehabilitation following neurological disease or injury. Adaptation, the initial stage of motor learning, involves short-term changes in motor performance in response to a new demand in the person’s environment. Repeated adaptation can improve skill performance and result in long-term skill retention. Locomotor adaptation is extensively studied using split-belt treadmill paradigms. In this study we explored whether bidirectional walking (BDW) on a split-belt treadmill can induce short-term gait adaptations. Twelve healthy volunteers participated in our single session, starting with 2 minutes of normal walking (NW), followed by four 5-minute blocks of BDW with a 1-minute passive rest in between blocks, and ending with another 2-minute of NW. We recorded body kinematics and ground reaction forces throughout the experiment. Participants quickly adapted to BDW with both legs showing decreased step lengths. However, only the backward-walking leg exhibited aftereffects upon returning to NW, indicating short-term adaptation. Notable kinematic changes were observed, particularly in hip extension and pelvis tilt, though these varied among participants. Our findings suggest that BDW induces unilateral adaptations despite bilateral changes in gait, offering new insights into locomotor control and spinal CPG organization.

**Key Points:**

- Locomotor adaptation extensively studied using split-belt treadmill paradigms with asymmetric belt speeds.
- This study examined whether bidirectional walking, i.e. walking with each leg stepping in opposite directions, would induce short-term adaptations in spatiotemporal characteristics of gait.
- Volunteers performed bidirectional walking with bilateral changes in step length.
- Despite equal but opposite belt speeds, volunteers exhibited a unilateral aftereffect of decreased step length in the leg which performed backwards stepping during bidirectional walking.

## Introduction

### Motor Learning

Humans can acquire and maintain motor skills throughout their lives. This ability arises from a diverse set of neurological processes collectively termed motor learning (for a review of motor learning taxonomy and terminology, see Krakauer et al. (2019)). It is motor learning that enables us to learn to draw a sketch, play a musical instrument, or ski down a snowy mountain. After neurological disease or injury (e.g. stroke, spinal cord injury) rehabilitative interventions rely on continued motor learning to enable recovery. One component of motor learning that has been extensively studied is adaptation, the short-term alteration in the execution of a previously acquired motor skill in response to change in sensory input (e.g., reaching within a force field) (Donchin et al., 2003; Scheidt et al., 2000; Shadmehr and Mussa-Ivaldi, 1994; Smith et al., 2006) or with a rotated field of view (Krakauer et al., 2000; Mazzoni and Krakauer, 2006; Peled and Karniel, 2012). Initially, these novel dynamics may impair performance (e.g., missing a desired reaching target). Repeated execution of the movement over several minutes changes movement execution and restores performance. If the altered sensory input is removed, the skill typically displays an aftereffect of decreased performance while the individual de-adapts to the perturbed dynamics. Repeated transient adaptations are believed to lead to more permanent, long-term skill acquisition (i.e., to persistent motor learning). The result is that adaptation to the altered sensory input occurs quickly, and the removal of this altered input produces little aftereffect. Thus, adaptation paradigms can be useful in studying motor learning.

### Locomotor Adaptation

Locomotor adaptation has been widely studied using split-belt treadmills (Reisman et al., 2005, 2007). Commonly, participants walk on the treadmill with two belts moving at different speeds. (Hagen et al., 2023; Torres-Oviedo and Bastian, 2012; Yokoyama et al., 2018; Malone and Bastian, 2014). This initially produces step-length asymmetry that is subsequently reduced by gait occurring independently in each limb (Choi and Bastian, 2007). This paradigm has been an invaluable tool for investigating locomotor adaptation. Furthermore, it has shown some promise as a therapeutic intervention for patients with asymmetries due to CNS injury or disease, such as stroke (Reisman et al., 2009, 2013; Frigon et al., 2015; Betschart et al., 2018).

### Bidirectional Walking

While many split-belt treadmill studies have explored speed differences between right and left legs, only a few have also studied bidirectional (i.e., hybrid) walking (Yang et al., 2005; Choi and Bastian, 2007). In bidirectional walking, the two treadmill belts move in opposite directions; one leg walks forward, the other walks backward. While not applicable in overground walking, bidirectional walking may be mechanically analogous to some turning strategies (Hase and Stein, 1999). This locomotor pattern decouples the stereotypical anti-phasic, flexor/extensor relationships of normal walking. During normal walking each leg alternates between extensor-driven swing phases and flexor-driven stance phases, and the two legs are out of phase with each other. In contrast, in bidirectional walking, the flexors drive the swing phase and the extensors drive the stance phase in the backward-walking limb. The ability of both infants (Yang et al., 2005) and adults (Choi and Bastian, 2007) to perform bidirectional walking suggests that the muscle pattern generation can be formed for each limb independently. Recent studies with decerebrated and spinalized cats (Lyakhovetskii et al., 2021; Audet et al., 2024) have demonstrated that bidirectional walking can be elicited with spinal stimulation. This suggests that, despite the functional decoupling of left and right limbs, spinal central pattern generators (CPGs) may be able to generate the muscle activity patterns for bidirectional walking which may provide insight into CPG organization (Veshchitskii et al., 2022). However, these previous studies focused on the feasibility of performing bidirectional walking or, in the case of Choi and Bastian (2007), whether left and right speed mismatches would independently induce adaptations. To the best of our knowledge, the occurance of short-term adaptation in bidirectional walking has not been directly tested. Nor whether this behavior is as stereotypical across individuals as normal locomotion.

### Study Approach and Objectives

The goal of the current study was to investigate whether bidirectional walking induces short-term adaptation and what strategies individuals use to perform it. We asked twelve volunteers to walk on a split-belt treadmill with the belts moving at equal speeds, but in opposite directions. We recorded body kinematics and treadmill force plate data for each participant during bidirectional walking and after returning to normal locomotion. We compared limb trajectories and joint angles between bidirectional and normal walking and evaluated whether bidirectional walking would induced aftereffects in spatiotemporal characteristics of gait. We hypothesized that (1) participants would alter their spatiotemporal gait characteristics, specifically double-stance time and stride length, and (2) these characteristics would demonstrate a transient aftereffect when returning to normal locomotion.

## Methods

### Ethical approval

All procedures were approved by the Institutional Review Board (IRB) of the Stratton VA Medical Center consistent with the standards of the *Declaration of Helsinki* (Protocol #: 1584762). Participants provided written informed consent to participate in the study.

### Participants

Twelve volunteers (mean age of 42.8 *±* 13.6 years) participated in the study. We included people who met the following criteria: no history of neurological deficits, ability to perform a 30-minute treadmill walk without assistance, and no prior experience with split-belt treadmill studies. Basic demographic data were collected (age, weight, height, and active hours per week, see Table 1). We also evaluated each participant’s foot dominance using the Waterloo Footedness Questionnaire. Because all our participants were found to have either mixed- or right-foot dominance, we conditioned the right leg to go in the opposite direction for everyone.

**Table 1:** Summary statistics of participant characteristics. Weight, height, age, and the preferred walking speed for normal locomotion are shown as the mean *±* the standard deviation across participants. The number of participants who were determined to be left-foot dominant (left dom.), right-foot dominant (right dom.), or mixed dominance (mixed dom.), as defined in the Waterloo Footedness Questionaire, are also provided.

| Participants | mean ± st. dev. |
| --- | --- |
| weight (kg) | $71.5 \pm 18.4$ |
| height (m) | $1.71 \pm 0.1$ |
| age (yrs) | $42.3 \pm 12.3$ |
| preferred speed (m/s) | $0.896 \pm 0.2$ |
| left dom. | 0 |
| mixed dom. | 5 |
| right dom. | 7 |

### Walking Paradigm

Each participant walked on a split-belt treadmill (Bertec Version 2.0; Columbus, OH) wearing a harness that was connected to the treadmill frame. The harness did not provide weight-bearing support unless the person fell. First, the participant’s preferred walking speed (PWS) was empirically determined by presenting five different speeds at (*±*0.2 m/s) increments. The speed was then kept at 80% of PWS throughout the session. Figure 1A outlines the walking paradigm explained here. Each session began and ended with 2 min of normal treadmill locomotion; the belts moved in the same direction at equal speeds, (i.e., normal locomotion(NL)/baseline). Then, over 1 min, right-belt speed was gradually decreased to zero and gradually increased to 80% PWS in the opposite direction, i.e., bidirectional locomotion bidirectional walking (BDW). The participant then performed four 5-min BDW training blocks with one min of quiet standing between the blocks.

**Figure 1:**
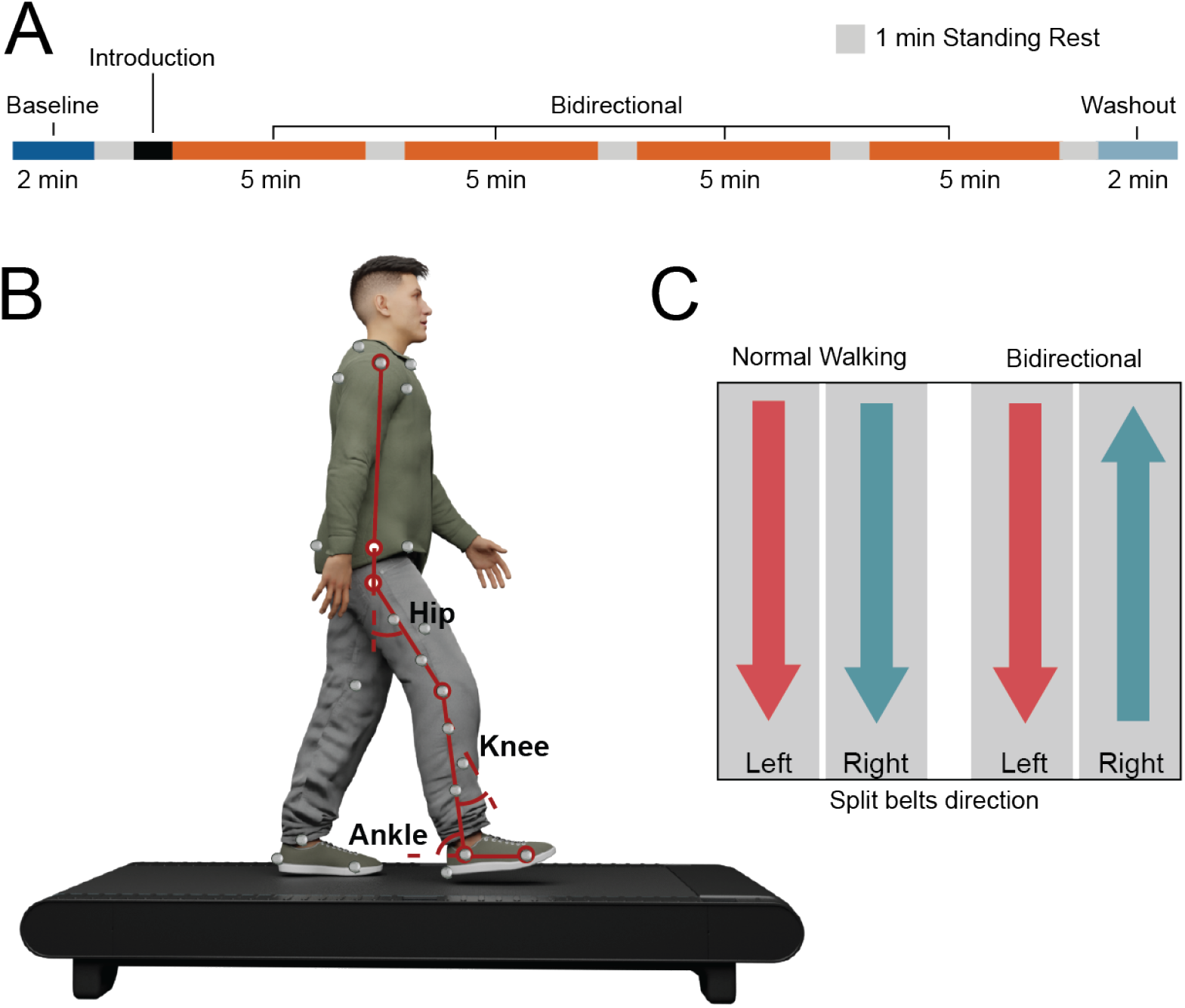
Experimental Paradigm. (A) Each participant was asked to perform the same walking paradigm. The session comprised six walking blocks. In the Baseline block, the participant walked with both belts moving in the same direction at 80% of their preferred walking speed. Next, after a brief 1-min introduction to walking with the belts moving at 80% preferred speed in opposite directions, i.e., right belt moving forward bidirectional walking (BDW), the participant performed BDW for four 5-min blocks. Finally, in the last block (Washout), they again performed normal locomotion. (B) Participant kinematics were recorded using 41 passive markers to track body kinematics. Belt directions for the left and right belts are shown in (C)

### Data Acquisition

Participants wore comfortable, form-fitting clothing, to facilitate accurate motion recordings. Loose-fitting clothing was secured using adhesive tape, as needed. Forty-one reflective markers were placed on bony landmarks of the head, torso and legs. Marker locations were based upon the Rajagopal musculoskeletal model which was used to perform inverse kinematics (Rajagopal et al., 2016). Three-dimensional marker locations were sampled at 100Hz using an 8-camera Qualysis motion capture system (Göteborg, Sweden). Prior beginning data collection, the participant was asked to hold a T-pose (arms abducted parallel to the ground with legs shoulder-width apart) for 10s. We used this marker positional data, and the participant’s height and weight, to facilitate scaling of a musculoskeletal model (see Kinematic Analysis).

Ground reaction forces and moments were recorded concurrently in three dimensions with the two force plates of the treadmill at a sampling frequency of 2000Hz. Prior to each session, the force plates were zeroed to ensure the precision and consistency of subsequent measurements.

### Data Processing

Data processing, analyses, and statistical analyses were performed using custom written scripts in Matlab (2023a) and R (R Core Team, 2020).

### Step Detection and Foot Positions

To define step onsets and offsets, we first applied a 2nd order, low-pass Butterworth filter (*F*_*c*_=20) to force plate measurements in the vertical (Z) direction to reject ambient electrical noise and other high frequency interference. We then determined step onsets and offsets for each leg by detecting threshold crossings of 50 N. We estimated foot positions during each step using the following equations (Corp, 2013):

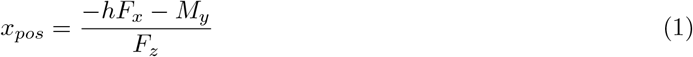

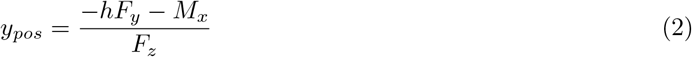

where *h* is the vertical distance from the treadmill surface to the force plate (0.015m), *F*_*x*_ is the force in the *x*-direction, *F*_*y*_ is the force in the *y*-direction, *M*_*x*_ is the moment around the *x*-direction, *M*_*y*_ is the moment around the *y*-direction, and *F*_*z*_ is the force in the *z*-direction.

### Kinematic Analysis

Joint angles were approximated using OpenSim 4.4 (Stanford University). We used a previously validated musculoskeletal model of the torso and lower limbs scaled to each participant’s anthropometry as shown in Figure 2A (Rajagopal et al., 2016). Scaling was performed with the OpenSim Scaling tool using marker locations collected at a static T-pose. Joint angles were approximated using the OpenSim InverseKinematics tool in accordance with data collection and simulation guidelines.

**Figure 2:**
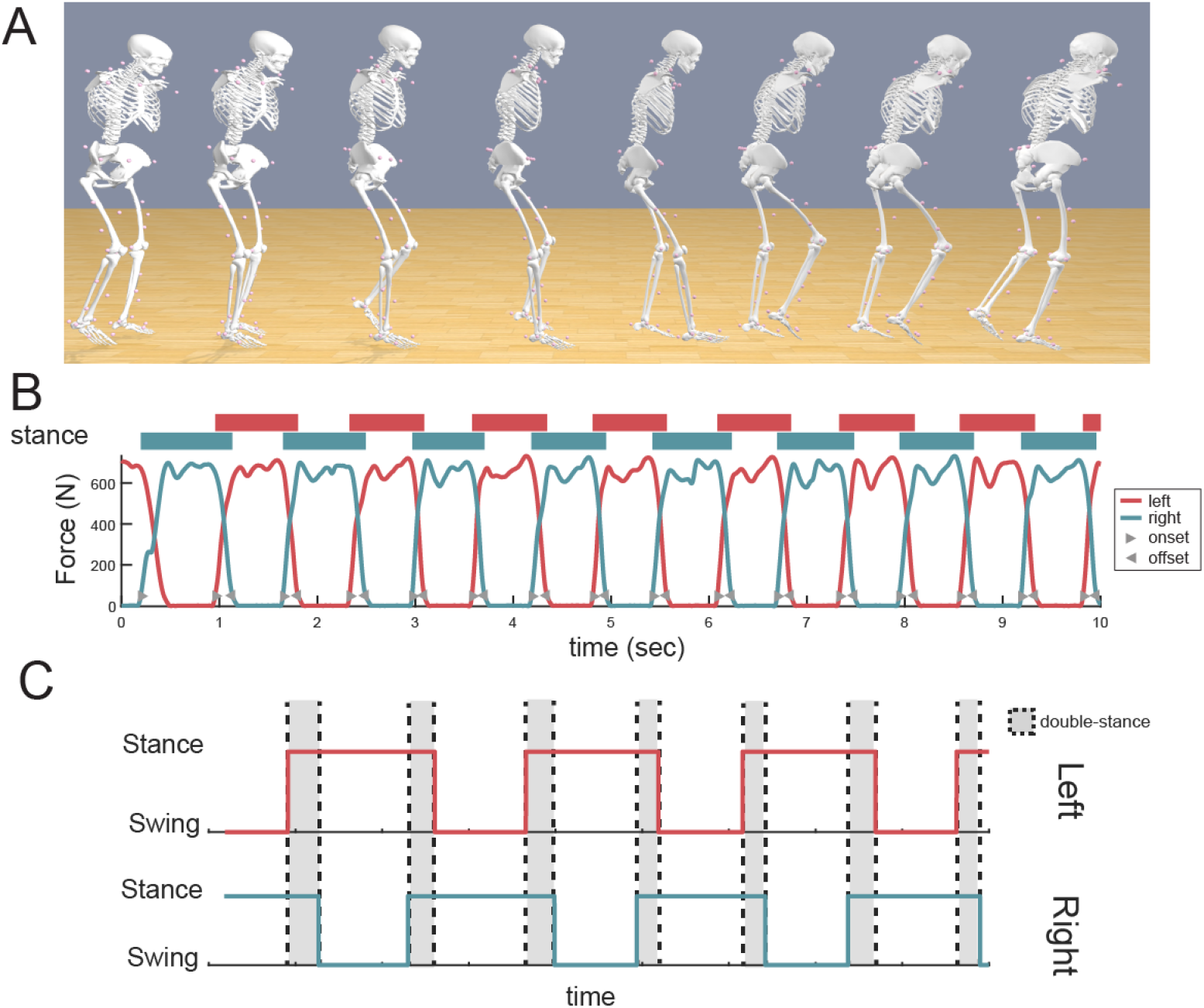
Data Processing. (A) To approximate joint angles from 3D marker positions we performed inverse kinematics using a musculoskeletal model of the torso and lower extremities (Rajagopal et al., 2016). (B) Step onsets and offsets were detected using force plate measurements from the treadmill. An illustrative example is shown with both the left force (red) and right force (blue) values shows along with the corresponding stance/swing phases. (C) Double-stance was calculated as the percentage of time that both legs were in stance phase divided by the total walking time.

### Double-stance

Using the detected onsets and offsets of each leg, we assigned each leg to be either in swing or stance (see Figure 2B). We then calculated double-stance as the percentage of time points where both legs were in stance divided by the total points in each block.

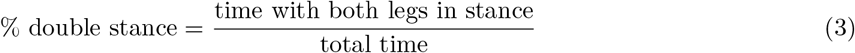

### Step Length

We defined step length as the distance between the left and right foot positions at the time of the stepping foot’s onset. We calculated the difference between foot positions in the anterior-posterior (Y) direction at the time of foot onset for the left and right feet independently.

## Statistical Analyses

To determine whether double stance time or step length differed between the baseline and washout phases, we calculated an average for each participant within each block. We then used a paired t-test (*α* = 0.05) to test the null hypothesis (*H*_0_) that double stance and step length did not differ between the baseline and washout phases. To determine whether these parameters adapted during the training phases, we similarly calculated an average for each participant within the training blocks. We fit a linear mixed effects model and calculated the 95% confidence interval for the overall model slope estimate.

## Results

### Kinematic changes

First we characterized how bidirectional walking differed from normal walking by analyzing limb trajectories and joint kinematics. Participants quickly adapted to the opposing belt directions – stepping forward with the left foot, while stepping backward with the right (See Figure 3A-C). Foot trajectories with the right leg appeared more variable during BDW than during NL as seen in Figure 3D-E. In addition, the vertical lift of right foot was smaller. While NL is characterized by repeated patterns of heel strike alternating between each leg (Figure 3G), BDW alternated between heel-strike for the left foot, followed by toe-strike for the right foot (Figure 3H).

**Figure 3:**
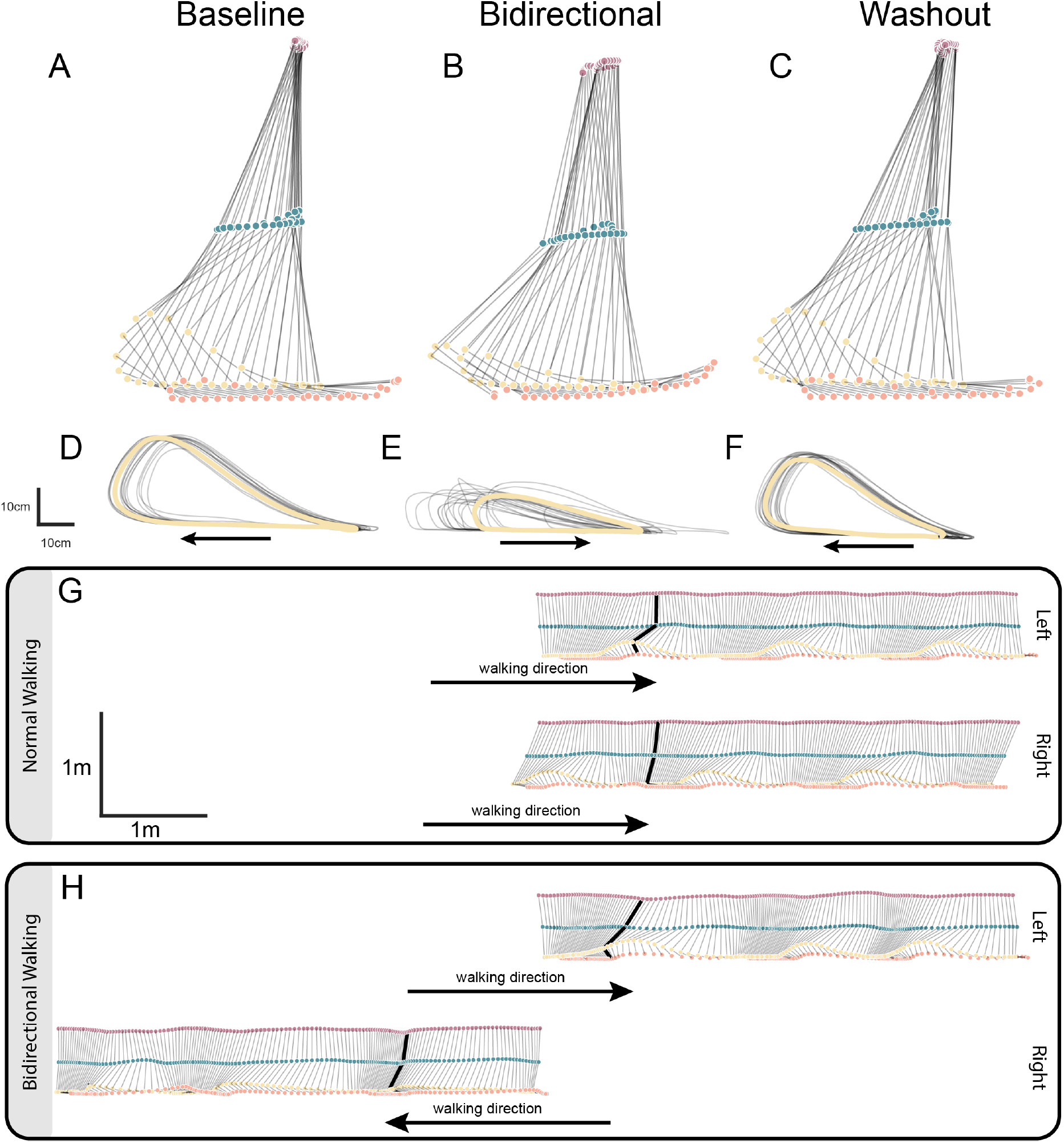
Leg Trajectories. A representative example of marker trajectories for the right leg are shown for the (A) baseline, (B) bidirectional, and (C) washout phases for a single gait cycle. (D, E, F) Mean heel trajectories for the same participant are shown in yellow for 20 steps shown in black. (G, H) Simulated gait patterns are shown for both normal and bidirectional locomotion for the right and left legs. Body translations are inferred based on the speed of the treadmill belts.

To better quantify gait characteristics, we calculated joint angles for both hips, knees, and ankles, along with pelvis orientation across all participants (Figure 4A). Joint kinematics for the left leg were largely unchanged between NL and BDW, although some subtle changes can be seen in knee and ankle flexion (e.g., Figure 4A, second to last row, rightmost two columns). Unsurprisingly, the right leg exhibits larger differences in joint angles with decreased hip extension and shifts in the phase of knee and ankle flexion. We also noticed marked changes in the pelvis tilt during the gait cycle (first row, first column in Figure 4). The combination of decreased hip extension with increased pelvis tilt resulted in a gait pattern analogous to an oscillating inverted pendulum (Figure 4B).

**Figure 4:**
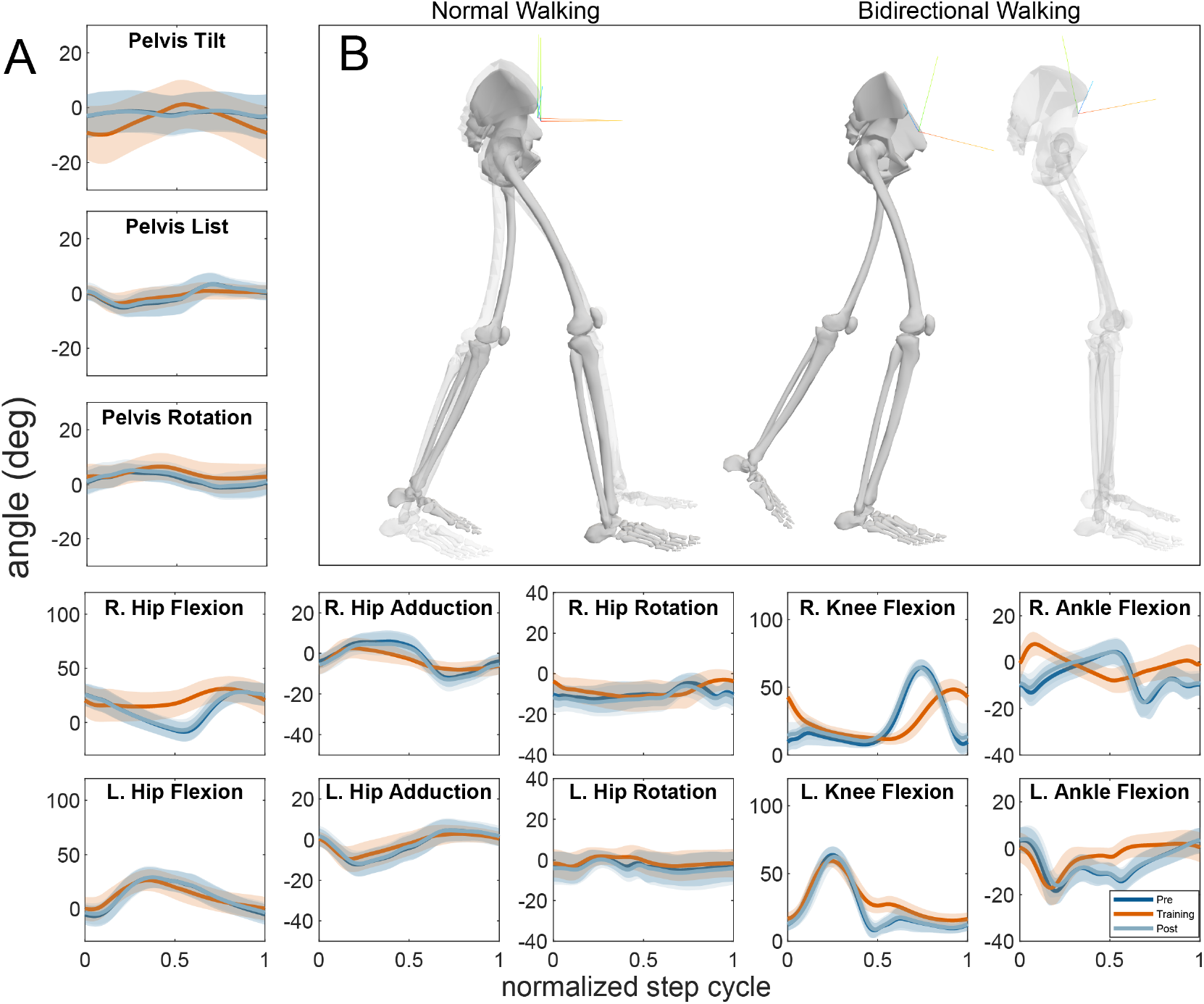
Joint Kinematics. (A)Joint angles for the hips, knees, and ankles along with pelvis orientation are shown for the baseline (dark blue), BDW (orange) and washout (light blue) phases. The y-axes have been scaled to the approximate joint range of motion for the hips, knees, and ankles. The solid line indicates the mean joint angle across participants and the shaded regions denote the standard deviation across participants. (B) Example of gait pattern for both normal and bidirectional walking. The opaque model shows the onset of right leg contact. The semi-transparent model shows the end of right leg stance.

### Double-stance time

We hypothesized that training with the BDW would change double-stance time and that this change would carry over to the NL washout block. We calculated percentage of the gait cycle in double-stance for each block using Eq. 3. Figure 5A shows the time in double-stance for each participant during each block. During NL, double-stance time ranged from 19.6-35.0% of the gait cycle, which is consistent with previous observations (Webster and Darter, 2019). We did not see any consistent changes in double-stance time across the baseline, BDW training, and washout blocks.

**Figure 5:**
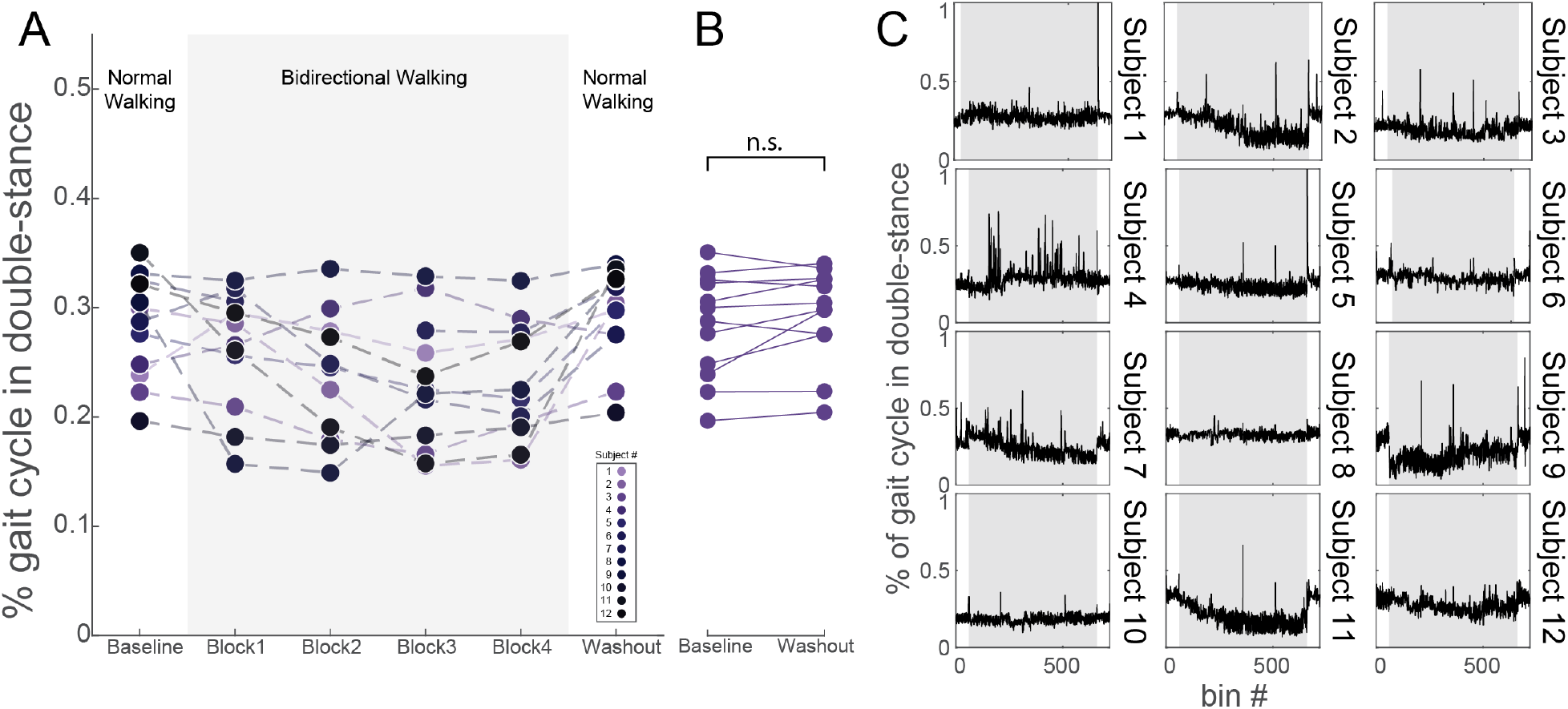
Double-stance. (A) The mean percentage of the gait cycle spent in double-stance is shown for each participant during each walking block. (B) Changes in the mean percentage of the gait cycle spent in double-stance is shown for each participant during the baseline and washout phases, i.e., before and after bidirectional training (C) The percentage of gait cycle in double-stance is shown for each participant over time. Each block was divided into 2 second bins and double-stance time was calculated for each bin.

We then tested the null hypothesis, (*H*_0_), that double-stance time did not differ between the baseline and washout blocks. We used a two-tailed, paired t-test with an *a priori* criterion of *α* = 0.05 and were unable to reject the null hypothesis (*t*_11_ = *−*1.74; *p* = 0.11; 95% CI: (*−*0.023, 0.003)) (Figure 3B). We then performed a *post hoc* subject-specific analysis to determine whether individual participants show changes in their double-stance time. We binned each participant’s session into 2-s time intervals and calculated the double-stance time for each bin (Figure 3C). Interestingly, some participants(2, 7, 9 & 11, 5C), show substantial decreases in their double-stance time during the training blocks, which disappeared when they returned to NL. In contrast, others (1, 5, 8, & 10, 5C) maintained stable double-stance time throughout all blocks.

### Step Length

Our second hypothesis was that training with the BDW would cause changes in step length and that this change would carry over to the NL washout block. We used foot positions calculated from the treadmill force plates to measure step length (see Figure 6A). To determine whether BDW training results in step length changes that carried over to NL, we calculated the difference between the left and right foot positions at the time of each step onset (Figure 2B). Here we defined a right step when the right limb is leading and visa versa. Step lengths were calculated independently for each leg because we did not assume that the step length, or changes in it, would be symmetrical during BDW.

**Figure 6:**
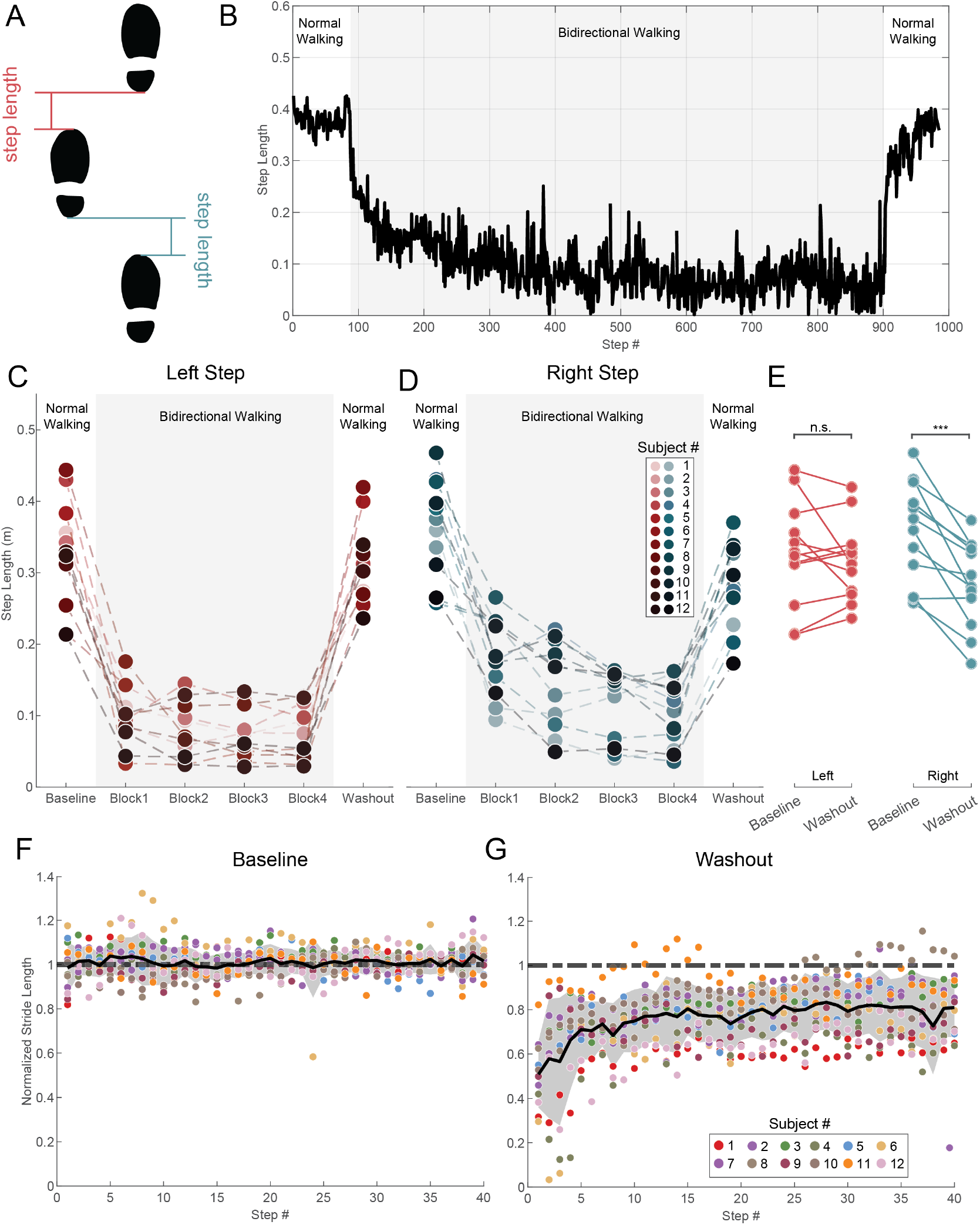
Step Length. (A) An illustrative depiction of step length.(B) A representative example of the step length for the right leg throughout an entire experimental session. The shaded region denotes steps performed during the training phases. (C,D) Mean step lengths for each participant are shown during each walking block for the left and right leg, respectively. (E) Changes in step length before and after the training phases are shown for the left and right leg. (F, G) The normalized step length for the last 40 steps of the baseline phase and the first 40 steps of the washout phase are shown. The different colored dots denote the value of individual steps for the 12 different participants. The solid black line indicates the mean across participants and the shaded region denotes *±* one standard deviation. The dotted line in (G) highlights the pre-training normalized value of 1.

Figure 6B shows a representative example of step length for the right leg across a session. The step length decreases substantially during BDW compared to NL, and lengthens after returning to NL. We calculated the mean step length for each participant in each block of the session for both legs (Figure 6, B&C). In both legs, the step length decreases during the BDW relative to pre-training NL. For the left leg, which is still walking forward, the decrease occurs in the first block and remains at a similar level throughout the remaining training blocks. Interestingly, the decrease in step length for the right leg, whose walking direction is reversed, increases gradually throughout the training blocks (Figure 6 B & C, shaded region). To confirm this observation, we used a linear mixed effects regression model, with random slopes and intercepts (i.e., within-participant intercept and slope) and overall slope and intercept fixed effects. To test the significance of the fixed slope effect, we used a likelihood ratio test. We found that slope of the right step size decreased significantly throughout the training blocks (estimate: *−*0.027; 95% CI (*−*0.039, *−*0.015); *p <* 0.001) while the slope of the left step size was not statistically different from 0 (estimate: *−*0.008*m*; 95% CI (*−*0.018, 0.002); *p* = 0.096.

We then tested the null hypothesis, (*H*_0_), that there is no change in average step length during NL performed prior to and after the BDW (baseline vs. washout) using a two-tailed, paired t-test (see Table 2). We rejected the null hypothesis for the right leg, but failed to reject it for the left (see Figure 6E & Table 2). In summary, our findings indicate that although step length decreases for both the right and left legs during the BDW, it is only the right leg decrease that transfers to NL.

**Table 2:** Two-tailed, paired t-test of mean step length before and after bidirectional walking.

|  | left step | right step |
| --- | --- | --- |
| $t_{11}$ | 0.761 | 5.172 |
| p-value | 0.463 | <**0.001\*** |
| mean | 0.010 | 0.069 |
| 95% CI | (−0.020, 0.040) | (0.041, 0.101) |

Next, we examined the time course of step length de-adaptation during the washout phase for the right leg. Figure 6F shows the step length at each right step for all participants during the baseline phase. Because individual participants walked at different speeds, each participant completed a variable number of steps in each 2-minute block. The first 40 steps are shown because all participants completed at least that number of steps. The step lengths have been normalized to each participant’s mean step length during this block to facilitate comparison across individuals. While step lengths vary step-by-step, most steps are within 1-2 standard deviations of the mean and remain relatively constant throughout the two-minute periods. Figure 6G shows the step-by-step step length during the washout phase. Here, we observed a decreased step length initially followed by a nonlinear increase induced by bidirectional training. After 2 minutes, most participants had still not recovered back to their pre-training step length.

The consistent decrease in step length across participants was surprising given the seemingly consistent kinematics during the baseline and washout phases in Figure 4. The relationship between step length and kinematics is deterministic so a decrease in step length would necessitate changes in the kinematics. We suspected that this contradiction may have arisen from averaging our kinematics across subjects so we performed a *post hoc* subject-specific kinematic analysis which examined joint angles on the time of right step onset, i.e., when the step length is decreased. We first used a univariate approach where we calculated the difference in joint angles during the washout phase with those during the baseline phase. Kinematic changes were highly variable across participants, e.g., while some participants achieved a shorter step length by increasing knee flexion, others did so by decreasing hip flexion. We then performed a Principal Component Analysis (PCA) and found that joint kinematics clustered separately for baseline and washout kinematics most participants. However here there was also high variability across people, suggesting that different people changed step length through different strategies (see Figure 7).

**Figure 7:**
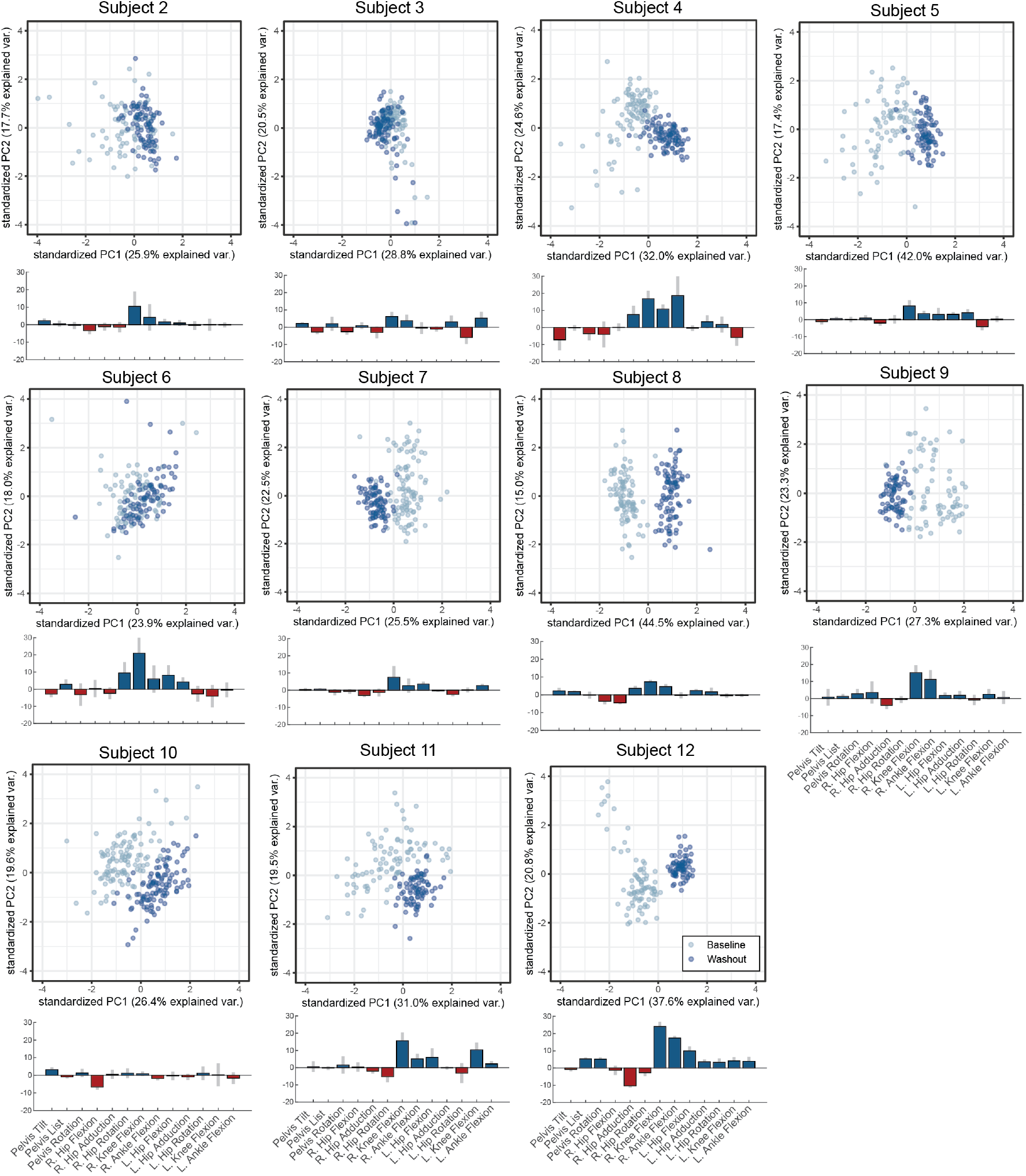
Changes in Joint Angle at Right Step. To examine what kinematic changes result in decreased step length during the washout phase, we performed a *post hoc*, subject-specific analysis of joint angles at the time of right foot onset. Participant 1 was excluded due to poor quality motion capture data. We calculated the mean difference in joint angles between early de-adaptation and late baseline (N=5 steps each). Increases in the de-adaptation phase are shown in blue, while decreases are shown in red for each participant. The grey bars indicate the 95% confidence interval of the mean. Most prominently, participants increased their right knee and ankle flexion to varying degrees. However, changes in other joints were more variable across participants, demonstrating individualized strategies to obtain and accommodate the decreased step length.

## Discussion

### Summary

In overground and tied-belt treadmill walking, the body is propelled forward by repeated, alternating patterns of limb stance and swing phases generated by similarly alternating recruitment of flexor and extensor muscles in each leg (Brown and Sherrington, 1997). Bidirectional walking–walking with both legs at equal speeds but in opposite directions–creates a challenge for locomotor control in that it also requires anti-phasic interlimb coordination but decouples the relative direction for each limb and the flexor/extensor contribution to swing and stance phases. This study shows that bidirectional walking differs from normal locomotion bilaterally – as shown by decreased left and right step lengths and increased pelvic motion – but causes short-term adaptation in only one limb as demonstrated by an aftereffect once returning to normal locomotion. These results demonstrate that, despite both legs moving at the same speed, bidirectional walking is an adaptive behavior which can be used in conjunction with other locomotor adaptation paradigms for studying shortterm adaptation and neural plasticity.

### Spatiotemporal Adaptation

Locomotor adaptation has been extensively studied using split-belt treadmill paradigms. Perhaps the most commonly used paradigm asks participants to perform forward walking with the belts moving at different speed ratios, e.g. 2:1, 3:1, etc. This paradigm imposes a spatiotemporal mismatch between the left and right limb. Initially, this induces a step length asymmetry in which the faster stepping leg displays a larger step size, compared to the leg on the slower moving belt. Over the course of 10-20 minutes, individuals use a combination of spatial and temporal modifications to decrease this asymmetry (Malone et al., 2012). Here, spatial modifications refer to changes in the relative foot placement of each limb. Changes in foot placement are achieved by changing the center of oscillation of one leg relative to the other. This effect can be visualized as a vertical shift in the slow-moving leg angle over time. Temporal modification refers to changes in the temporal phase of each leg relative to the other or a horizontal shift in one leg angle over time relative to the other. These spatiotemporal changes occur independently for each limb (Choi and Bastian, 2007); they are the result of error-driven feedback due to inter-limb step length asymmetry caused by the mismatched speed.

In our study, bidirectional walking induced asymmetry not due to differing speeds – both belts moved at the same speed – but stepping in opposite directions. Similar to paradigms using asymmetric speeds, participants unilaterally adapted their step lengths, as shown by a transient aftereffect upon returning to normal locomotion. However, these results are unique from previous studies in that unilateral adaptation was induced by a gait pattern that required a bilateral change in motor strategy. Both the forward and backward walking limbs displayed a decreased step length during bidirectional walking, but only the backward walking limb showed aftereffects due to adaptation. These results demonstrate two key findings:

(1) unilateral adaptations can carry over from one locomotor pattern to a second pattern that differs enough that it requires bilateral changes to perform it;
(2) while adaptations may occur independently in each limb, the control strategy for each limb is interdependent, i.e. the movement of left limb in bidirectional walking is not equivalent to performing forward walking.

### Implications of Bidirectional Walking on CPG Organization

Thomas Graham Brown demonstrated that locomotor rhythms could be generated in the absence of cortical and afferent input over a century ago and proposed that rhythmogenesis was generated by two half-center oscillators in the spinal cord that maintained interlimb coordination (Brown and Sherrington, 1997; Brown, 1914). Since Brown’s seminal work, numerous theoretical and computational models of the spinal CPG have expanded or adapted Brown’s half-center model, but maintain that interlimb coordination is preserved through intercommissural spinal connections between motor neurons of the left and right limb (Székely et al., 1969; Duysens and Pearson, 1976; Perret and Cabelguen, 1980; Grillner et al., 1981; Burke et al., 2001; Yakovenko et al., 2005; Brownstone and Wilson, 2008; McCrea and Rybak, 2008; Yakovenko et al., 2018). Logically, these models were developed to explain unidirectional – usually forward – walking. However, recent studies in decerebrated and spinalized cats have shown that bidirectional walking can also be performed without higher brain centers or afferent feedback (Lyakhovetskii et al., 2021; Audet et al., 2024). These findings suggest that spinal CPG architecture is sufficient for rhythmogenesis and muscle activity formation in both bidirectional and normal walking. Furthermore, the ability of the feline CPG to maintain interlimb rhythm despite the limbs moving in opposite directions is consistent with higher-order CPG models, which separate rhythm generation and muscle pattern formation into separate interneuron layers (Veshchitskii et al., 2022).

In our study, participants obviously performed bidirectional walking with the added input of higher centers and afferent feedback. Nevertheless, there are some notable differences in their performance compared to decerebrate and spinalized cats that may provide insight into the role of this additional input. First, the kinematics of each limb in the decerebrate cats resembled the normal kinematics of forward and backward walking when performing bidirectional walking, i.e., the “forward walking” limb resembled normal forward walking and the “backward walking” limb resembled normal backward walking(Lyakhovetskii et al., 2021). In constrast, in our study, the participants changed the step lengths of both limbs during bidirectional walking, choosing to shorten both while allowing their pelvis to oscillate along the anterior-posterior axis. It is possible that these differing strategies may be a byproduct of differences in the experimental paradigm. For example, the cats’ pelvises were stereotactically fixed to provide weight support and thus unable to oscillate. Alternatively, these differences may be the result of an explicit change in locomotor strategy which required supraspinal input to execute. Furthermore, we observed unilateral aftereffects in the backward walking limb. While unilateral adaptations are commonly reported at different speed ratios – including during bidirectional walking (Choi and Bastian, 2007) – here the belts are moving at the same speed but in opposing directions. Thus, while bidirectional walking may be executed with only CPG architecture, in intact humans repeated performance causes short-term, unilateral adaptations.

## Conclusions

In conclusion, we investigated whether bidirectional walking would induce aftereffects in spatiotemporal parameters of gait. We found that during bidirectional walking the participants changed their step length bilaterally but displayed unilateral aftereffects indicative of short-term adaptation. Changes in kinematics and double-stance were more varied across participants. This study shows that locomotor adaptations may be induced by strategies other than differing limb speeds and offers a complimentary paradigm to do so.

## Additional Information

### Data Availability

Data supporting the results of the study is available upon reasonable request.

### Author Contributions

All experiments were performed at the Samuel S. Stratton VA Medical Center, Albany, NY. The authors Russell L. Hardesty (RH), Helia Motjabavi (HM), Darren E. Gemoets (DG), and Jonathan R. Wolpaw (JW) approve the accuracy and integrity of the study. The contributions of the each author are listed below.

- Conceptualization/Experimental Design: RH, HM, JW
- Data Collection: RH, HM
- Data Analysis: RH, DG
- Writing/Manuscript Preparation: RH, HM, DG JW

## Acknowledgements

The authors would like to acknowledge Drs. Sergiy Yakovenko and Jonathan Carp for their valuable scientific input. We would also like to acknowledge the National Center for Adaptive Neurotechnologies (NCAN) and the Samuel S. Stratton VA Medical Center for administrative, logistical support and use of their facilities. Drs. Wolpaw and Gemoets are employees of the Samuel S. Stratton VA Medical Center. The contents of this manuscript do not represent the views of the US Department of Veterans Affairs or the United States Government.

